# A joint probabilistic model of human scene and object recognition via non-hierarchical residual computation

**DOI:** 10.1101/2025.05.02.651866

**Authors:** Kosuke Nishida, Isamu Motoyoshi

**Affiliations:** Kanazawa University: Kanazawa Daigaku; The University of Tokyo

**Keywords:** scene perception, object recognition, peripheral vision, coarse-to-fine, variational autoencoder, figure-ground segmentation

## Abstract

Visual object and scene recognition have been extensively studied, but separately. We here propose that the two processes could be intrinsically linked in the neural system. We developed a Joint Residual Variational Autoencoder (JRVAE) with two networks: VAE1 for coarse scene recognition and VAE2 for object recognition using residuals from VAE1’s reconstructions. Our model demonstrates emergent functional specialization when conditioned on information reduction in peripheral vision, with quantitative analysis confirming VAE1 excels at the representation of scenes while VAE2 specializes in that of objects. This architecture naturally implements figure-ground segmentation and aligns with neurobiological evidence of distinct cortical pathways. Our findings suggest residual computation enables joint visual processing that mirrors human perception’s coarse-to-fine principle in perception.

## Introduction

By looking at natural images, humans can easily recognize the overall scene and individual objects in it. The neural mechanisms underlying object and scene recognition have been extensively studied across the fields of psychophysics, neurophysiology, and computer vision (DiCarlo et al., 2012; Epstein, 2005; Epstein & Baker, 2019; Kriegeskorte et al., 2008; Tanaka, 1996). However, object and scene recognition mechanisms, while sometimes compared to each other, have been considered separate topics and studied independently. In particular, many studies of object recognition have focused on the perception of objects placed against a uniform field or within a very limited size of contextual background (Cichy et al., 2014; Cohen et al., 2017; Grill-Spector et al., 2001; Wolfe et al., 2011). Yet, the real visual field is far broader, requiring the visual system to detect and select objects from complex natural environments. Therefore, in realistic contexts, object recognition is inseparable from scene recognition, and both should be understood in an integrated manner. The present study aims to capture the relationship between scene and object recognition from this unified perspective.

Traditionally, the cognitive processes involved in separating objects from natural scenes have been thought to rely on low-level information processing, such as local contrast of visual features (Itti & Koch, 2000; Koch & Ullman, 1985) or concentration of top-down attention based on prior knowledge (Desimone & Duncan, 1995; Reynolds & Chelazzi, 2004). For example, models that guide attention based on visual saliency (Bruce & Tsotsos, 2009; Itti & Koch, 2000; Kummerer et al., 2017) represent typical theories of this approach. However, it is well known that the visual system can grasp the meaning of an entire scene in a very short time, with humans able to accurately identify scene categories within a short latency of approximately 100 milliseconds after stimulus presentation (Li et al., 2002; Thorpe et al., 1996). The process of separating objects from scenes, accompanied by such rapid recognition, cannot be explained merely by the accumulation of feature contrast processing and is likely based on more comprehensive scene understanding. Indeed, models based on saliency maps have shown limitations in predicting human gaze positions within natural scenes (Castelhano et al., 2009; Henderson et al., 2007; Henderson & Hayes, 2018). Therefore, the standpoint that object recognition is performed based on relatively high-order statistical properties of background scenes provides a more reasonable explanation of actual visual processing (Henderson, 2017; Whitney & Yamanashi Leib, 2018).

A large part of the scene comprises peripheral vision. Neural representations in peripheral vision have lower spatial resolution compared to central vision, with information represented in a more summary statistical form. In fact, the spatial sampling rate of the retina decreases rapidly with increasing eccentricity (Curcio & Allen, 1990), and peripherally viewed images synthesized based on low-level image statistics (so-called metamers) are difficult to visually distinguish from the original images (Freeman & Simoncelli, 2011). These statistics have been shown to correlate with neural representations in visual area V2 (Freeman et al., 2013), supporting the notion that peripheral vision comprehends scenes through summary statistics. In real visual environments, it is difficult to clearly extract the contours of unknown objects appearing in peripheral vision, which is consistent with the aforementioned physiological findings. Nevertheless, conventional object detection models, such as those for visual search(Wolfe, 1994, 2021; Wolfe & Horowitz, 2017), have not adequately considered these characteristics of peripheral vision. On the other hand, recent models based on statistical representations of peripheral vision have been shown to effectively explain human performance in various tasks, including visual search and classification (Rosenholtz et al., 2012), analogous to surface statistics (e.g., Motoyoshi et al., 2007). Additionally, the visual system appears capable of extracting semantic information about scenes with considerable accuracy using only the reduced information obtained from peripheral vision. Indeed, studies using statistically synthesized images of scenes have reported that scene recognition performance is not significantly impaired compared to original images (Ehinger & Rosenholtz, 2016).

Based on these findings, object detection can be understood as a process of separating individual objects, in some way, from coarse scene representations across the entire visual field. In fact, the essential information of a scene is known to be extracted rapidly, prior to object recognition (Oliva, 2005; Oliva & Torralba, 2006), and this sequential order of processing is consistent with the “coarse-to-fine” principle in vision(Hegdé, 2008; Watt, 1987). This principle has been widely observed in various classical psychophysical tasks, such as binocular stereopsis(Y. Chen & Qian, 2004) and hierarchical pattern discrimination(Navon, 1977).

The ventromedial visual pathway, which governs scene recognition (Grill-Spector & Weiner, 2014; Rolls, 2024), has the characteristic of prioritizing statistical information in the peripheral visual field (Baldassano et al., 2016; Silson et al., 2015; Silson, Groen, et al., 2016; Silson, Steel, et al., 2016), with cortical areas showing selective responses to scenes (PPA, MPA, OPA) being representative examples (Choo & Walther, 2016; Epstein & Baker, 2019; Ganaden et al., 2013; Marchette et al., 2015; Mégevand et al., 2014; S. Park et al., 2011). Features of these areas include processing superiority from peripheral vision and selective responses to textures (J. Park & Park, 2017). In contrast, the ventrolateral visual pathway, responsible for object recognition, prioritizes central vision and does not adequately represent statistical information in peripheral vision. Indeed, many neurons belonging to visual areas IT/TE selective for objects (Hung et al., 2005; Rust & DiCarlo, 2010; Yao et al., 2023) have receptive field centers biased toward central vision (DiCarlo & Maunsell, 2003). This differentiation of visual pathways could enable a processing flow from coarse scene comprehension by the ventromedial pathway to detailed object identification by the ventrolateral pathway, facilitating the “coarse-to-fine” visual information processing. This processing is thought to be achieved through feedback projections in information flow from scenes to objects (Brandman & Peelen, 2017; de Lange et al., 2018; Peelen et al., 2024), which is consistent with Hochstein’s reverse hierarchy theory (Hochstein & Ahissar, 2002). Additionally, studies using backward masking, EEG, and fMRI (Groen et al., 2018; Scholte et al., 2008; Seijdel et al., 2021) have suggested the existence of segmentation mechanisms through recurrent connections within the ventral visual pathway.

As described above, scene and object recognition, while based on different visual pathways and processing stages, are suggested to be closely interconnected in actual visual cognition. Particularly in real visual environments constrained by peripheral vision, the integrated processing of coarse scene information and detailed object information is particularly important. In spite of this, computational models that simultaneously implement and integrate this dual-processing architecture and appropriate feedback mechanisms within an unsupervised learning framework are notably absent in the current literature. Here, in the same way as object recognition has been explained within an unsupervised learning framework such as β-VAE models (Higgins et al., 2021), one can predict that even for scenes, which are generally considered more complex than objects, a similar unsupervised method would apply by taking into account the information reduction in peripheral vision during scene recognition. This research aims to fill the above-mentioned gap of scene-object recognition by attempting to construct an integrated visual cognition model that utilizes the statistical structure of natural images under unsupervised learning while considering the characteristics of peripheral vision and coarse-to-fine information transfer.

## Results

### 1. Model and Dataset Preparation

While the ventral visual pathway has been extensively modeled as a hierarchical object recognition network (DiCarlo et al., 2012; Yamins et al., 2014), the computational principles underlying scene recognition and its relationship to object processing remain underexplored. We implemented them as deep generative models using variational autoencoders (Kingma & Welling, 2014). In fact, during visual working memory, item-specific signals in the primary visual cortex (V1) are confined to deep and superficial layers, avoiding the middle layer, while such signals span all cortical depths in V2 and V3, highlighting distinct laminar processing of top-down versus bottom-up information (Lawrence et al., 2018).

Our Joint Residual Variational Autoencoder (JRVAE) framework proposes that scene processing occurs prior to and facilitates object recognition through a residual computation mechanism. This arrangement functionally mirrors the observed anatomical distinctions between ventromedial (scene) and ventrolateral (object) processing pathways (Rolls, 2024).

To test this hypothesis, we developed and trained two complementary networks:

- VAE1: Trained to learn scene representations
- VAE2: Trained on the residuals between inputs and VAE1’s reconstructions We utilized two datasets:

1. Scene-only dataset: Consisting of 1,090 images representing 6 distinct scene categories viewed from multiple angles
2. Scene-object dataset: Comprising 7,047 images with 6 scene categories and 6 object categories positioned centrally

To simulate the peripheral bias observed in scene-selective regions, we applied Gaussian blur to peripheral regions of input images. VAE1 was initially trained on the scene-only dataset for 150 epochs, then briefly exposed (2 epochs) to the scene-object dataset. The residuals between the scene-object inputs and VAE1’s reconstructions were computed, center-cropped to 1/4 size, and fed to VAE2 for 20 epochs of training. All models were enabled with microssacade mode that augments the input following a Gaussian distribution (mu=center, sigma=3.0, maximal shift=7.0) to improve model stability. In VAE2, “temporal average smoother” and “activation threshold” were introduced to denoise the temporally inconsistent residuals as well as amplifying temporally consistent residuals. These are detailed in Appendix. Both VAE1 and VAE2 are beta-VAEs (Burgess et al., 2018). VAE1’s beta was fixed to 1.0 and VAE2’s beta was fixed to 4e-2 during training.

### 2. Emergent Figure-Ground Segmentation Through Residual Computation

As predicted by our computational framework, VAE1 effectively reconstructed peripheral elements while leaving central regions (containing objects) as residuals (Figure 2). This process naturally implemented figure-ground segmentation without explicit supervision or attention mechanisms. The residual computation rendered peripheral regions mostly gray in the input to VAE2, indicating that VAE1 had successfully predicted these scene elements. While we only consider the case where the object is already in the center, this could also cope with the cases where objects are in the periphery (see “soft-max policy” in Appendix).

**Figure 1.**
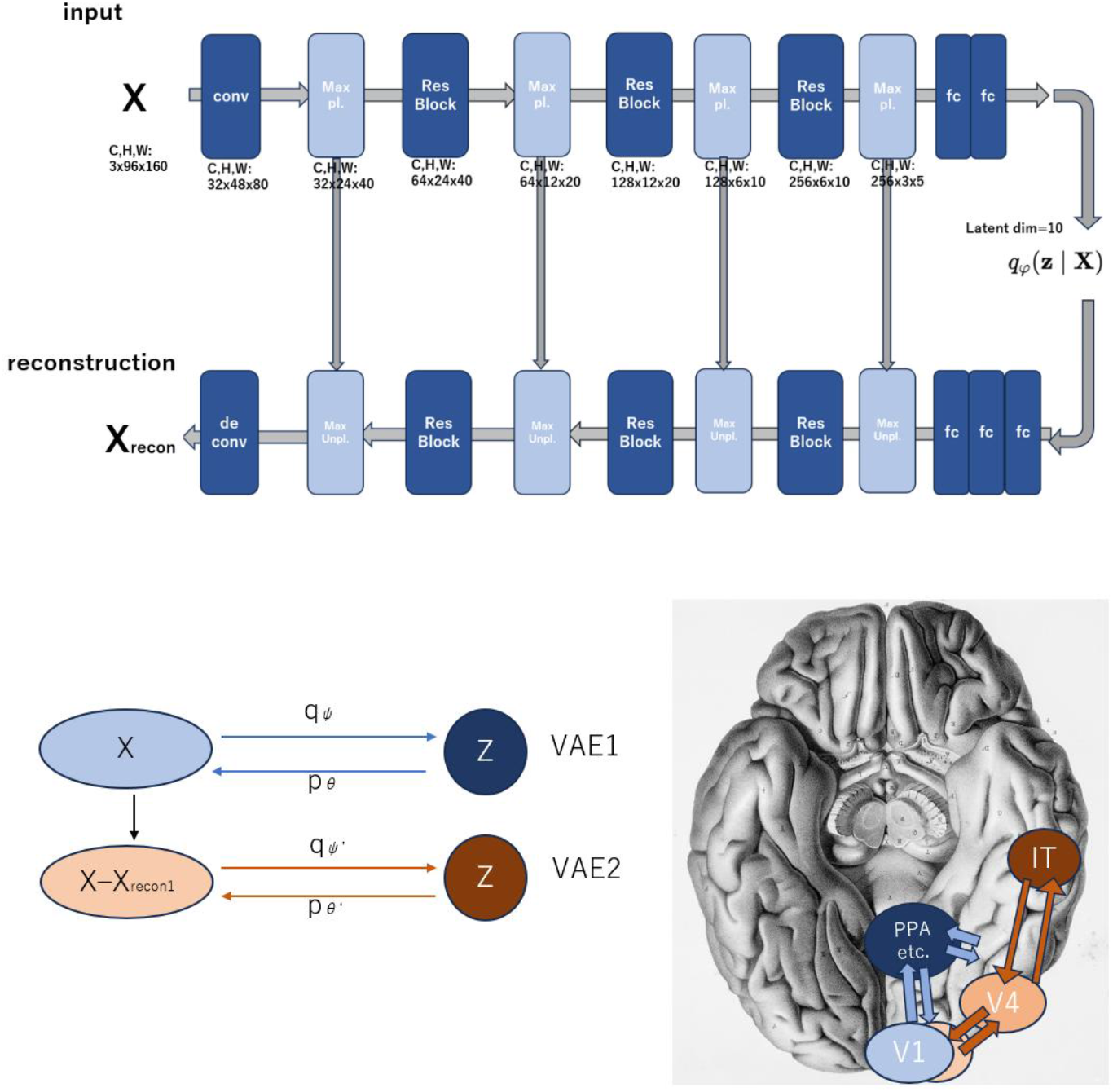
(top) the architecture of a VAE used in our model. The input has size (C, H, W) = (3, 96, 160). (bottom)schematic architecture of our proposed model (Joint Residual VAEs). (bottom left) The input images will be fed into VAE1, which produces the reconstructions. The “residuals” between the input and the reconstruction will then be fed into VAE2. The input to VAE2 is smoothed by “temporal average smoother” and “activation threshold”, which we will detail in Appendix. Our hypothesis is that VAE1 corresponds to scene perception and VAE2 performs object recognition. (bottom right) Blue arrows indicate ventromedial pathways, while brown arrows show ventrolateral pathways as revealed by Rolls, 2024. The inferior view of a brain image is given by Foville, licensed under CC BY 4.0, via Wikimedia Commons (<u>File:”Traite complet de l’anatomie…”,Foville, 1844 Wellcome L0019135.jpg - Wikimedia Commons</u>).

**Figure 2.**
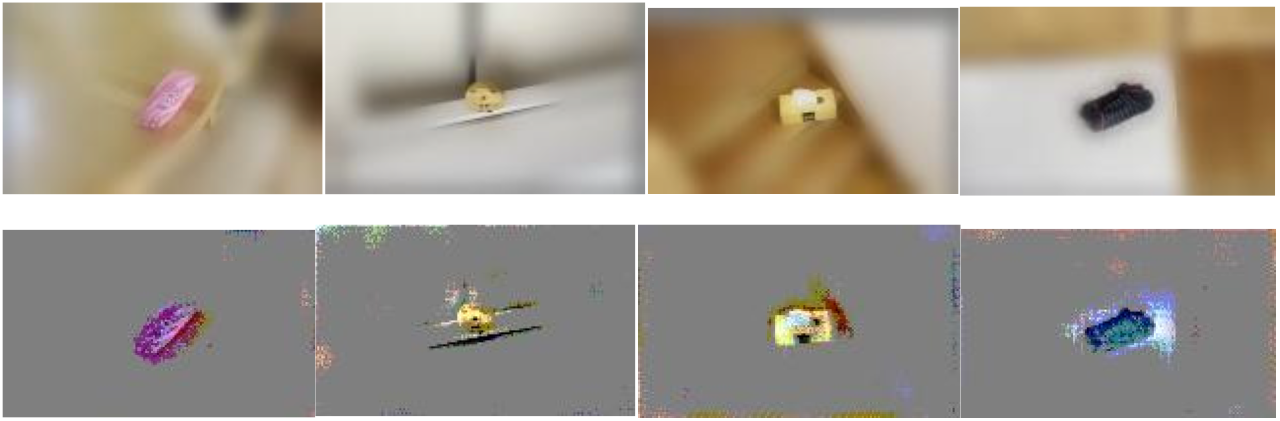
Examples of VAE1’s input(top) and VAE2’s input(bottom). The actual input (above) is substracted from VAE1’s reconstruction, which results in the VAE2’s input(bottom).

### 3. Functional Specialization Emerges Through Network Interdependency

To analyze the distribution of latent representations in our trained models, we applied Uniform Manifold Approximation and Projection, or UMAP (McInnes et al., 2018) to visualize the 10-dimensional latent space in two dimensions. This dimensionality reduction technique preserved the topological structure of the high-dimensional manifold while projecting the latent vectors of our test set (n=764), which comprised images of 6 distinct scene categories and 6 object categories. UMAP visualization enabled qualitative assessment of clustering patterns that emerged in the latent space of each network.

Dimensionality reduction via UMAP revealed that VAE1 learned to effectively cluster scene categories while showing minimal object categorization capability (Figure 3, top panels). Conversely, VAE2 demonstrated strong object clustering with minimal scene representation (Figure 3, middle panels).

**Figure 3.**
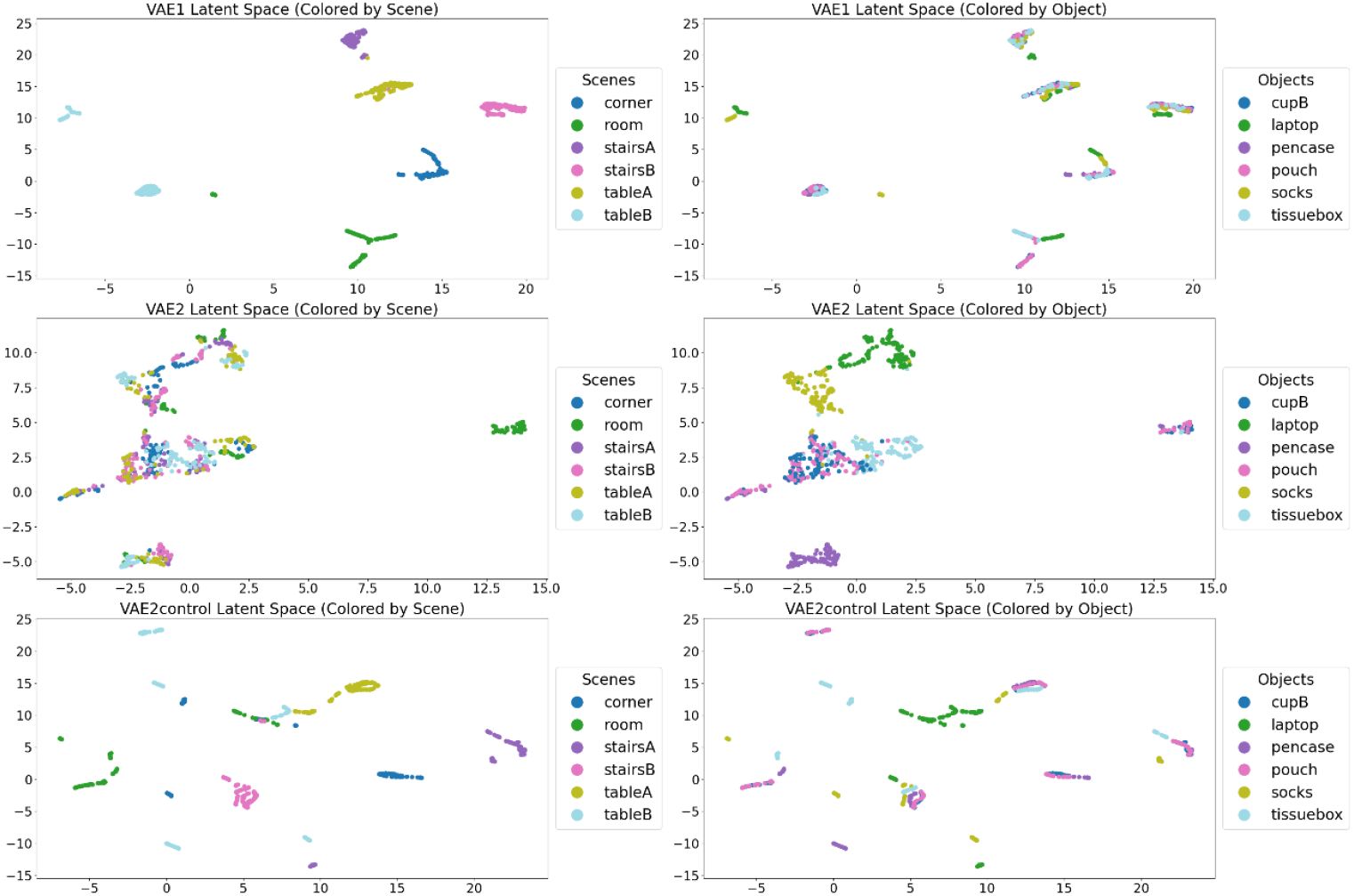
UMAP visualization of VAE1(top left and top right), VAE2(middle left and middle right), and VAE2_control (bottom left and bottom right). These dots are latent variables under the test set(n=764). The left column shows latent variable clusters given scene labels, whereas the right column indicates latent variable clusters with object labels.

**Figure 4.**
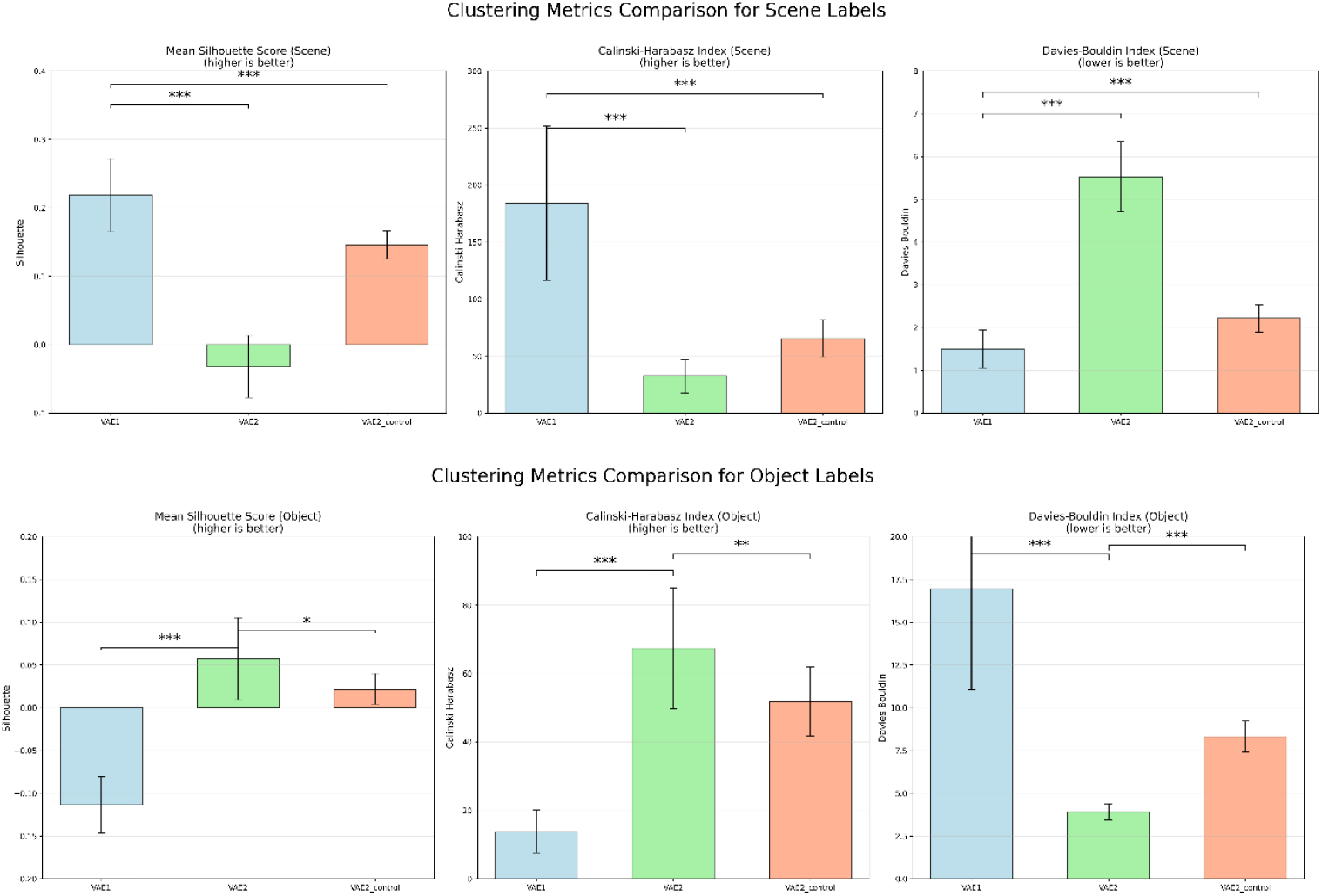
Silhouette score, Calinski-Harabasz index and Davies-Bouldin index for scene labels(top) and object labels(bottom). For scene labels, scores were compared between (VAE1 and VAE2) or (VAE1 and VAE2 control). For object labels, scores were compared between (VAE2 and VAE1) or (VAE2 and VAE2 control).

Crucially, this functional specialization depended on the residual computation between networks. When VAE2 was trained directly on the scene-object dataset without VAE1’s preprocessing (i.e., VAE2_control condition), it defaulted to scene classification despite having identical architecture and objective function (Figure 3, bottom panels). This demonstrates that unsupervised networks can develop complementary specializations through structured interdependency.

### 4. Quantitative Assessment of Representational Quality

To quantitatively evaluate the representational quality of each network, we applied three established clustering metrics to the latent embeddings: **1. Silhouette Score**: Measures how similar objects are to their assigned cluster compared to other clusters (higher is better) **2. Calinski-Harabasz Index**: Evaluates cluster separation based on the ratio of between-cluster to within-cluster dispersion (higher is better) **3. Davies-Bouldin Index**: Quantifies the average similarity between clusters (lower is better). These metrics are frequently used to evaluate clustering performance (Arbelaitz et al., 2013).

Analysis was done by python sklearn.metric in Python.VAE2control was prepared to confirm that the difference in VAE2’s performance was not caused solely by image clipping and differences in beta (between VAE1 and VAE2) defined in loss functions.

That is, VAE2control’s input is (C, W, H) = (3, 80, 48) clipped so that the object is placed in the center and beta’s value in the loss function is set 4e-2.

All pairwise comparisons between models revealed statistically significant differences (*** p < 0.001; ** p<0.01; * p<0.1) under two-sided Welch’s t-test. Here, fifteen models(N=15) were prepared for each VAE (VAE1, VAE2, VAE2_conrol) initialized with different random seeds. In scene representation performance, VAE1 demonstrated superior scene clustering the highest Silhouette Score (0.218) and Calinski-Harabasz Index (184.016), and the lowest Davies-Bouldin Index (1.498). VAE2 performed poorly on scene classification (Silhouette: -0.032; Calinski-Harabasz: 32.444; Davies-Bouldin: 5.530). VAE2_control showed intermediate performance (Silhouette: 0.146; Calinski-Harabasz: 65.430; Davies-Bouldin: 2.219). In object representation performance, VAE2 excelled at object clustering with a positive Silhouette Score (0.057), high Calinski-Harabasz Index (67.388), and low Davies-Bouldin Index (3.925). VAE1 performed poorly on object classification (Silhouette: -0.114; Calinski-Hrabasz: 13.772; Davies-Bouldin: 16.918). VAE2_control again showed intermediate performance (Silhouette: 0.022; Calinski-Harabasz: 51.858; Davies-Bouldin: 8.321).

These findings suggest that the dual-network JRVAE architecture can spontaneously develop specialized representations for scenes and objects through unsupervised learning, with scene processing facilitating object recognition through residual computation, aligned with the UMAP visualization result in the previous section.

## Discussion

We focused on a simple yet critical setting where the selection of an object is needed, that is, images of 6 kinds of objects embedded in 6 kinds of scenes, viewed from different angles, with peripheral vision blurred. Our results demonstrate that the Joint Residual Variational Autoencoder (JRVAE) can effectively specialize in scene and object recognition through distinct but complementary networks under such conditions.

Through unsupervised learning, VAE1 developed specialized representations for scenes while VAE2 effectively clustered objects through residual computation, as demonstrated by our UMAP visualizations and quantitative clustering metrics (Silhouette Score, Calinski-Harabasz Index, and Davies-Bouldin Index). This functional specialization only emerged with the residual connection between networks, as suggested by the control condition where VAE2 showed an intermediate performance when trained directly on the scene-object dataset. This computational framework addresses the three key challenges identified in our introduction: the precedence of scene perception over object recognition, the fundamental limitations of peripheral vision, and the feedback mechanisms from scene to object processing networks.

Since our architecture is based on beta-VAE, this can be seen as an extension of a previous object recognition model (Higgins et al., 2021) to both scene perception and object recognition, conditioned on limitations in peripheral vision. JRVAE is comprised of two VAEs interconnected through the input layer. If VAEs or similar models are actually implemented in the brain (Csikor et al., 2023; Jiang & Rao, 2024; Marino, 2021; Nagy et al., 2020; Summerfield & De Lange, 2014; Vafaii et al., 2023), it is possible that the relationship between scene and object recognition could also be implemented in the form of JRVAE. Aside from beta-VAEs simply trying to understand the object, there is also an attempt to understand the entire scene in the field of machine learning—compositional scene representation learning (Burgess et al., 2019; Locatello et al., 2020).These models output an “attention mask” to segment an object, often via VAE, eventually comprising the entire scene. JRVAE may look similar to these in the sense that the network tries to clip the input to subsequently recognize an object, resulting in both representing scenes and objects. However, JRVAE goes opposite to this direction: JRVAE’s generative model of a scene itself acts like a mask, following the “coarse-to-fine” scheme in humans. As we demonstrated, this is made possible by the dual architecture, each of which simply performs variational inference. Other approaches, such as contrastive learning methods, are very much common for the past several years for object recognition in the ventrolateral pathway. It is also known that these models do predict the explained variance of IT or TE neural activities very well (Zhuang et al., 2021). Nevertheless, because conventional contrastive learning methods do not have a generative model or any architecture for augmenting the input image itself, it is still not clear how such an architecture can implement some form of figure-ground segmentation as well as scene perception without a generative architecture. Future investigation might have to systematically explore the integration of contrastive and generative approaches to address this potential limitation.

Importantly, the JRVAE framework may extend beyond the scene-object relationship to encompass other hierarchical visual categories. Just as objects emerge as residuals from scene predictions, finer visual features ---such as texts in a book, a door mirror of a car or a specific pattern of a single flower petal---might emerge as residuals from object predictions of a book, a car or a flower, etc. This suggests the possibility of a third VAE in our architecture, specializing in processing “finer visual features” that deviates from typical object statistics. This extension seems in line with known functional specialization in the ventrolateral prefrontal cortex (VLPFC), which shows selectivity for specific visual features beyond object identity (Everling et al., 2006; Hussar & Pasternak, 2009; Mendoza-Halliday & Martinez-Trujillo, 2017), and has been associated with “feature-based attention (Bichot et al., 2019; Martinez-Trujillo, 2022).” What has been labeled as “feature-based attention” in the VLPFC might actually represent this third level of residual processing in our framework. Similarly, the frontal eye field’s (FEF) role as “spatial attention (Gregoriou et al., 2012; Wardak et al., 2006),” together with superior colliculus (SC), could be reinterpreted as implementing saccade planning toward locations based on simple image statistics such as high residual signals-essentially, directing the fovea toward visual information not predicted by current scene or object models in a way similar to what we demonstrated as “soft-max policy” in Appendix. This reconceptualization might offer a hint about the mechanism of attentional shifts, and invites rigorous empirical investigations.

Moreover, from a computational perspective, if each network in the JRVAE can distinguish N categories, then a system composed of M such networks would effectively perform N^M categorization from a single input. However, this combinatorial increase in representational capacity would come with constraints that parallel human perception. The cascading nature of residual computation necessitates progressively smaller inputs, resulting in diminished capacity to represent peripheral visual information—a limitation that might correspond to phenomena like change blindness in human perception(Rensink et al., 1997; Simons & Levin, 1997).

It is also worth noting that scene recognition shares deep connections with spatial navigation (Epstein & Baker, 2019; Marchette et al., 2015). While scene perception has been primarily studied in visual processing pathways, spatial navigation is largely attributed to the hippocampal-entorhinal system (Stachenfeld et al., 2017). The Clone Structured Causal Graph model (George et al., 2021), a prominent model of the hippocampal-entorhinal system, naturally incorporates the contextual considerations— which first-order Markov models generally fail to account for— in spatial navigation by allowing “clones” for each latent variable. Surprisingly, CSCG shows response patterns quite similar to those observed in place cells, grid cells and many other (Raju et al., 2024). Although our current JRVAE implementation does not fully address contextual facilitation (Bar, 2004)—the enhanced recognition of objects when presented in congruent scenes versus neutral backgrounds—incorporating CSCG-like structures could potentially resolve this limitation, as well as incorporating actions. Contextual facilitation of objects has been confirmed through fMRI decoding studies(Brandman & Peelen, 2017) as well as that of emotions(Z. Chen & Whitney, 2019). Similarity between spatial navigation and episodic recall of objects has been pointed out (Buzsáki & Moser, 2013), suggesting that scene and object classifiers may have a common computational principle for temporal sequences.

Our research not only invites reconsiderations of scene-object relationships and attention mechanisms in humans, but also might potentially present an opportunity to bridge computational models of the hippocampal-entorhinal system with those of visual processing. By reformulating visual cognition via non-hierarchical residual computation, we provide a much simpler understanding of the complementary yet distinct processes that seem to underlie human visual system.

A limitation of our current implementation is its scalability to complex, diverse natural images. While our results demonstrate proof-of-concept for the residual computation principle, the VAE architecture employed in this study would likely struggle with datasets of the complexity and diversity of ImageNet or similar collections. Future work should explore more robust generative architectures that could better scale to naturalistic complexity while preserving the core residual computation mechanism.

Furthermore, our experimental paradigm tested only scenarios with a single, centrally located object, whereas natural visual scenes typically contain multiple objects with varying spatial relationships. Future research must address how the residual computation scales to multi-object settings, potentially through iterative or recursive applications of the residual mechanism.

## Code and data availability

The full code will be available no later than the publication date.

## Author contributions

Conceptualization: KN, IM.

Data curation: KN.

Formal analysis: KN.

Investigation: KN, IM.

Methodology: KN, IM.

Software: KN.

Supervision: IM.

Visualization: KN.

Writing – original draft: KN, IM.

Writing – review & editing: KN, IM.

## Acknowledgement

This study was supported by JSPS KAKENHI JP23K25751 and JP24H01540 to IM.

## Appendix

### Hyperparameters

In our research, hyperparameters of the model were as follows:

Latent_dim=10

Learning_rate1=1e-5

Learning_rate2=1e-4

Beta1=1.0

Beta2=4e-2

Epoch1=150

Epoch2=20

Notably, setting β_2_=4e-2 (beta value in VAE2’s loss function) or near proved critical for model performance. Here, we stopped training when epoch=20, and in all cases, random seed was fixed to 42. Higher values (β_2_≥0.5) consistently resulted in posterior collapse (Fig. S1). While we acknowledge the potential impact of latent dimensionality and learning rate parameters, comprehensive hyperparameter optimization falls outside the scope of this investigation.

**Figure S1.**
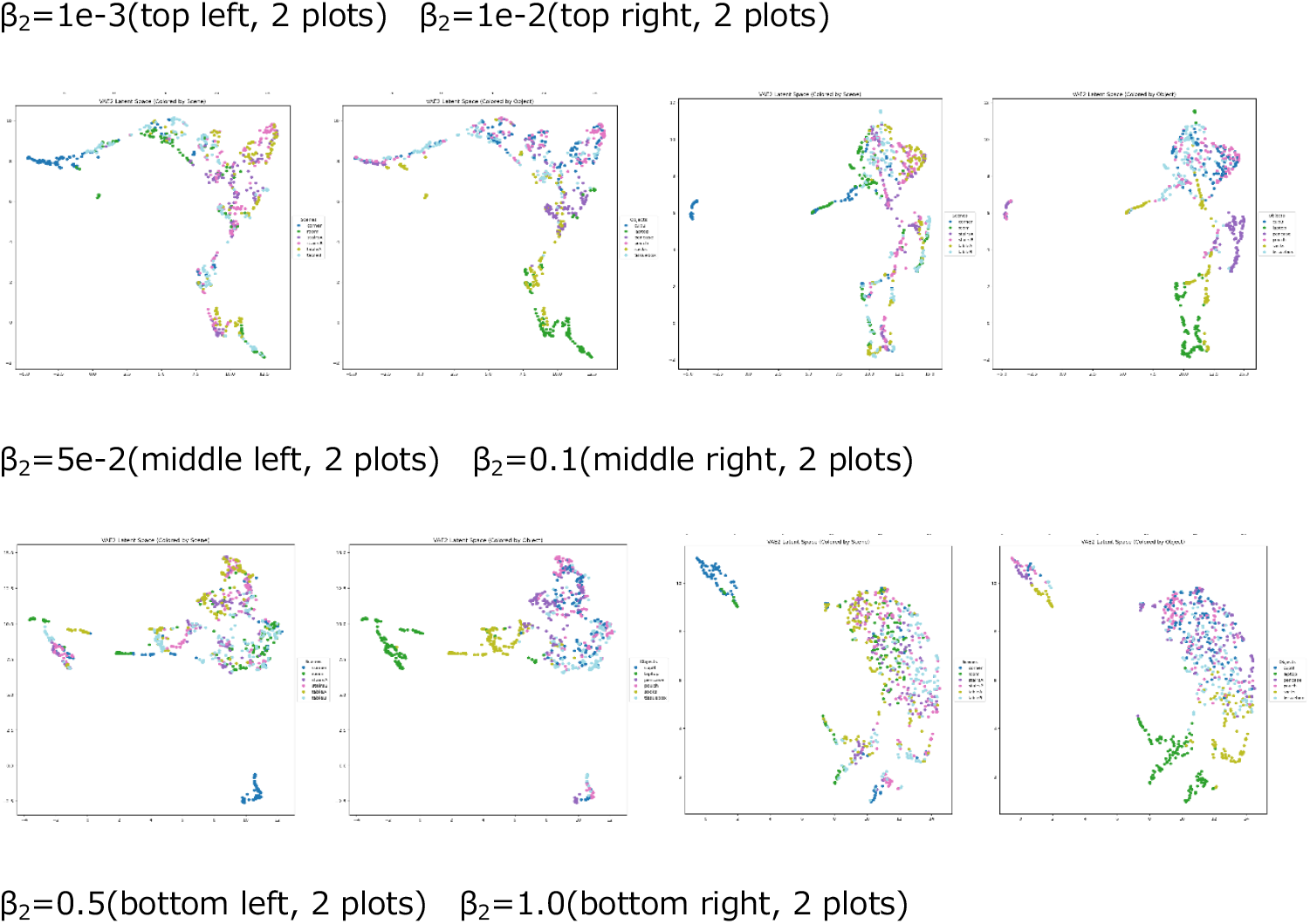
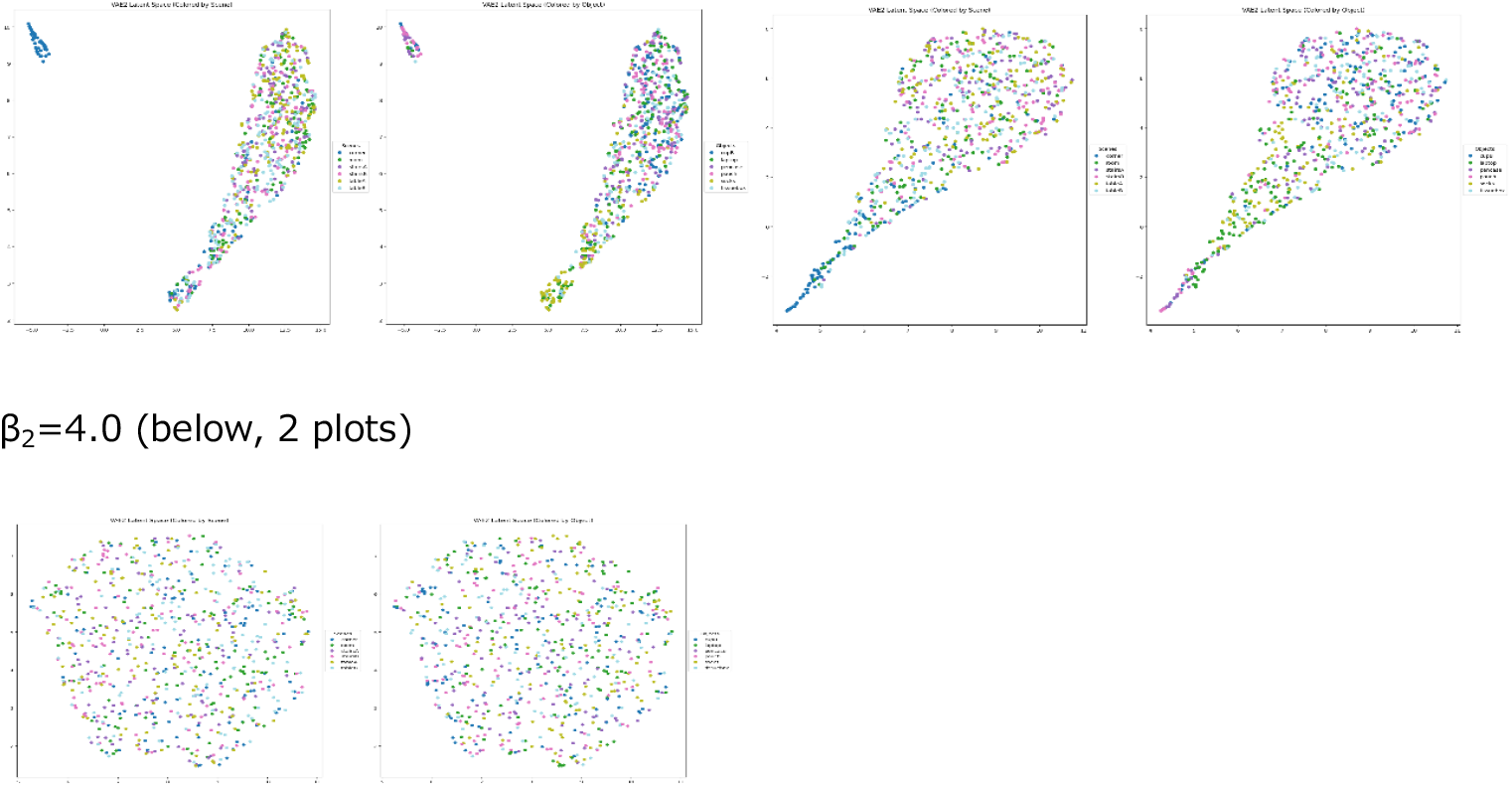
UMAP visualization of VAE2 latent space when epoch=20 with different beta2 values.

### soft-max policy

In the case where residuals are in the periphery (here, a bee is placed in a scene-only dataset), one would have to apply an augmentation such that it moves the residuals to the center. As an example, consider a position vector ***a***_***ij***_ (i = 1, 2, … ; j = 1, 2, …), where (i, j) indicates a position in the image. Let’s say this vector is sampled following a distribution π:

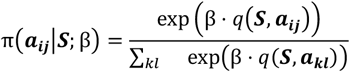

where q(***S, a***_***ij***_) = |*S*_*ij*_ | + *C* (*C* is an arbitrary constant and ***S*** is the input). *S*_*ij*_ is simply the pixel value ([-1, 1]) in the position (i, j), and q is the utility function. That is, a vector ***a***_***ij***_ gets a high weight if it has a high contrast relative to pixel values in other positions. This resembles “soft-max policy” in reinforcement learning but we already define q in advance for simplicity, which is the fundamentally different part from typical reinforcement learning, where it generally only defines the reward and the state-value function and action-value function are estimated by the agent). If we consider the expectation of this vector as ***A***, conditioned on ***S*** (input) we get

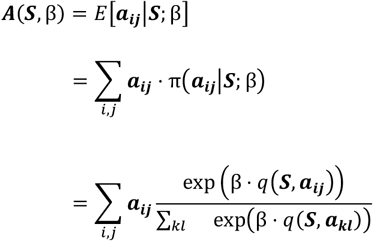

Increasing β from 1 to 25 results in (fig). blue point indicates β=1(start point) and the red point is β=25(end point). we are not claiming that this is what neuronal populations in FEF and SC are exactly doing in the form of saccadic planning. This is just one example of placing “high residual signals” to the center with no background based on a populational approach.

Here, VAE1’s output was clipped in the range [-1,1] using torch.clamp(), which is also the case in the models stated in the result section.

**Figure S2.**
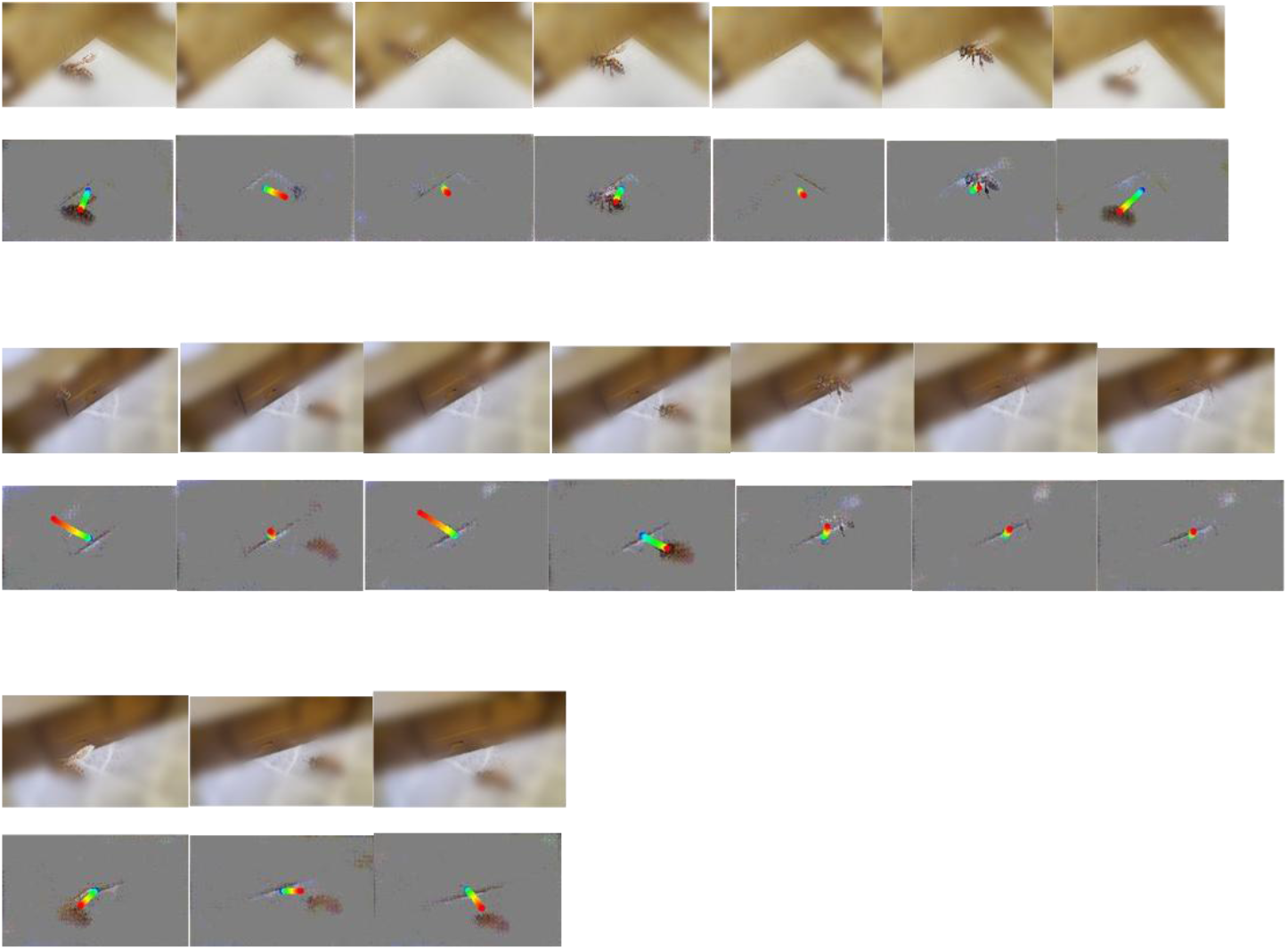
(top row) indicates examples of a new dataset (scene-only dataset with a bee added, position randomized), wheras (bottom row) shows the residuals between the actual input and VAE1 prediction that was learnt on scene-only dataset.

### Temporal Smoothing

Temporal smoothing is implemented in the compute_category_averages method of AveragedResidualDataset to stabilize the residual representations across consecutive frames by averaging them. This technique helps reduce noise and emphasize consistent patterns in the visual information that persists over time. Window-based Averaging: The implementation uses a sliding window approach with a configurable window size (default = 10 frames). Sliding Window Progression: The window slides forward one frame at a time, creating a temporal smoothing effect where each output frame represents the average of window_size input frames. Category-based Processing: The smoothing is applied separately to each category of scenes (e.g., “corner_pencase”), preserving category-specific visual characteristics.

### Activation Threshold

The activation threshold mechanism implements a form of sparse coding by selectively keeping only the most significant residual values and suppressing noise. This is implemented through the thresholding operation in the compute_category_averages method. Threshold Parameter: The mask_scale parameter (default = 0.20) determines the activation threshold. Binary Masking: The operation creates a binary mask where only residual values with absolute magnitude greater than mask_scale are preserved (set to 1), while smaller values are suppressed (set to 0).

### Amplification Mode

Amplification mode enhances the contrast of the thresholded residuals to make salient features more pronounced. This is implemented after the thresholding operation in the compute_category_averages method. Scaling Factor: The amp_scale parameter (default = 1.5) controls the amplification strength. Range Preservation: The torch.clamp function ensures that values remain within the valid range of [-1, 1] after amplification, preventing saturation artifacts. Contrast Enhancement: Multiplying by amp_scale increases the magnitude of the preserved residual values, making subtle features more prominent.

### Microsaccade Mode

Microsaccade mode simulates the small, involuntary eye movements that occur during human fixation. Gaussian Distribution: The shifts follow a Gaussian (normal) distribution, centered at zero with a Magnitude Limiting: The shifts are clipped to a maximum value to prevent excessive displacement. Image Shifting: The implementation uses padding and cropping operations to shift the image by the calculated offsets.

## Notes

### Competing Interest Statement

The authors have declared no competing interest.

## References

Arbelaitz, O., Gurrutxaga, I., Muguerza, J., Pérez, J. M., & Perona, I. (2013). An extensive comparative study of cluster validity indices. Pattern Recognition, 46(1). 10.1016/j.patcog.2012.07.021

Baldassano, C., Esteva, A., Fei-Fei, L., & Beck, D. M. (2016). Two distinct Scene-Processing networks connecting vision and memory. ENeuro, 3(5). 10.1523/ENEURO.0178-16.2016

Bar, M. (2004). Visual objects in context. In Nature Reviews Neuroscience (Vol. 5, Issue 8). 10.1038/nrn1476

Bichot, N. P., Xu, R., Ghadooshahy, A., Williams, M. L., & Desimone, R. (2019). The role of prefrontal cortex in the control of feature attention in area V4. Nature Communications, 10(1). 10.1038/s41467-019-13761-7

Brandman, T., & Peelen, M. V. (2017). Interaction between scene and object processing revealed by human fMRI and MEG decoding. Journal of Neuroscience, 37(32). 10.1523/JNEUROSCI.0582-17.2017

Bruce, N. D. B., & Tsotsos, J. K. (2009). Saliency, attention and visual search: An information theoretic approach. Journal of Vision, 9(3). 10.1167/9.3.5

Burgess, C. P., Higgins, I., Pal, A., Matthey, L., Watters, N., Desjardins, G., & Lerchner, A. (2018). Understanding disentangling in \${\textbackslash}beta\$-{VAE}. 1804.03599 [Cs, Stat].

Burgess, C. P., Matthey, L., Watters, N., Kabra, R., Higgins, I., Botvinick, M., & Lerchner, A. (2019). MONet: Unsupervised Scene Decomposition and Representation. https://arxiv.org/pdf/1901.11390

Buzsáki, G., & Moser, E. I. (2013). Memory, navigation and theta rhythm in the hippocampal-entorhinal system. In Nature Neuroscience (Vol. 16, Issue 2). 10.1038/nn.3304

Castelhano, M. S., Mack, M. L., & Henderson, J. M. (2009). Viewing task influences eye movement control during active scene perception. Journal of Vision, 9(3). 10.1167/9.3.6

Chen, Y., & Qian, N. (2004). A coarse-to-fine disparity energy model with both phase-shift and position-shift receptive field mechanisms. Neural Computation, 16(8). 10.1162/089976604774201596

Chen, Z., & Whitney, D. (2019). Tracking the affective state of unseen persons. Proceedings of the National Academy of Sciences of the United States of America, 116(15). 10.1073/pnas.1812250116

Choo, H., & Walther, D. B. (2016). Contour junctions underlie neural representations of scene categories in high-level human visual cortex: Contour junctions underlie neural representations of scenes. NeuroImage, 135. 10.1016/j.neuroimage.2016.04.021

Cichy, R. M., Pantazis, D., & Oliva, A. (2014). Resolving human object recognition in space and time. Nature Neuroscience, 17(3). 10.1038/nn.3635

Cohen, M. A., Alvarez, G. A., Nakayama, K., & Konkle, T. (2017). Visual search for object categories is predicted by the representational architecture of high-level visual cortex. Journal of Neurophysiology, 117(1). 10.1152/jn.00569.2016

Csikor, F., Meszéna, B., & Orbán, G. (2023). Top-down perceptual inference shaping the activity of early visual cortex. BioRxiv.

Curcio, C. A., & Allen, K. A. (1990). Topography of ganglion cells in human retina. Journal of Comparative Neurology, 300(1). 10.1002/cne.903000103

de Lange, F. P., Heilbron, M., & Kok, P. (2018). How Do Expectations Shape Perception? In Trends in Cognitive Sciences (Vol. 22, Issue 9). 10.1016/j.tics.2018.06.002

Desimone, R., & Duncan, J. (1995). Neural mechanisms of selective visual attention. In Annual Review of Neuroscience (Vol. 18). 10.1146/annurev.ne.18.030195.001205

DiCarlo, J. J., & Maunsell, J. H. R. (2003). Anterior inferotemporal neurons of monkeys engaged in object recognition can be highly sensitive to object retinal position. Journal of Neurophysiology, 89(6). 10.1152/jn.00358.2002

DiCarlo, J. J., Zoccolan, D., & Rust, N. C. (2012). How does the brain solve visual object recognition? In Neuron (Vol. 73, Issue 3). 10.1016/j.neuron.2012.01.010

Ehinger, K. A., & Rosenholtz, R. (2016). A general account of peripheral encoding also predicts scene perception performance. Journal of Vision, 16(2). 10.1167/16.2.13

Epstein, R. A. (2005). The cortical basis of visual scene processing. In Visual Cognition (Vol. 12, Issue 6). 10.1080/13506280444000607

Epstein, R. A., & Baker, C. I. (2019). Scene Perception in the Human Brain. In Annual Review of Vision Science (Vol. 5). 10.1146/annurev-vision-091718-014809

Everling, S., Tinsley, C. J., Gaffan, D., & Duncan, J. (2006). Selective representation of task-relevant objects and locations in the monkey prefrontal cortex. European Journal of Neuroscience, 23(8). 10.1111/j.1460-9568.2006.04736.x

Freeman, J., & Simoncelli, E. P. (2011). Metamers of the ventral stream. Nature Neuroscience, 14(9). 10.1038/nn.2889

Freeman, J., Ziemba, C. M., Heeger, D. J., Simoncelli, E. P., & Movshon, J. A. (2013). A functional and perceptual signature of the second visual area in primates. Nature Neuroscience, 16(7). 10.1038/nn.3402

Ganaden, R. E., Mullin, C. R., & Steeves, J. K. E. (2013). Transcranial magnetic stimulation to the transverse occipital sulcus affects scene but not object processing. Journal of Cognitive Neuroscience, 25(6). 10.1162/jocn_a_00372

George, D., Rikhye, R. V., Gothoskar, N., Guntupalli, J. S., Dedieu, A., & Lázaro-Gredilla, M. (2021). Clone-structured graph representations enable flexible learning and vicarious evaluation of cognitive maps. Nature Communications, 12(1). 10.1038/s41467-021-22559-5

Gregoriou, G. G., Gotts, S. J., & Desimone, R. (2012). Cell-type-specific synchronization of neural activity in FEF with V4 during attention. Neuron, 73(3). 10.1016/j.neuron.2011.12.019

Grill-Spector, K., Kourtzi, Z., & Kanwisher, N. (2001). The lateral occipital complex and its role in object recognition. Vision Research, 41(10–11). 10.1016/S0042-6989(01)00073-6

Grill-Spector, K., & Weiner, K. S. (2014). The functional architecture of the ventral temporal cortex and its role in categorization. In Nature Reviews Neuroscience (Vol. 15, Issue 8). 10.1038/nrn3747

Groen, I. I. A., Jahfari, S., Seijdel, N., Ghebreab, S., Lamme, V. A. F., & Scholte, H. S. (2018). Scene complexity modulates degree of feedback activity during object detection in natural scenes. PLoS Computational Biology, 14(12). 10.1371/journal.pcbi.1006690

Hegdé, J. (2008). Time course of visual perception: Coarse-to-fine processing and beyond. In Progress in Neurobiology (Vol. 84, Issue 4). 10.1016/j.pneurobio.2007.09.001

Henderson, J. M. (2017). Gaze Control as Prediction. In Trends in Cognitive Sciences (Vol. 21, Issue 1). 10.1016/j.tics.2016.11.003

Henderson, J. M., Brockmole, J. R., Castelhano, M. S., & Mack, M. (2007). Visual saliency does not account for eye movements during visual search in real-world scenes. In Eye Movements: A Window on Mind and Brain. 10.1016/B978-008044980-7/50027-6

Henderson, J. M., & Hayes, T. R. (2018). Meaning guides attention in real-world scene images: Evidence from eye movements and meaning maps. Journal of Vision, 18(6). 10.1167/18.6.10

Higgins, I., Chang, L., Langston, V., Hassabis, D., Summerfield, C., Tsao, D., & Botvinick, M. (2021). Unsupervised deep learning identifies semantic disentanglement in single inferotemporal face patch neurons. Nature Communications, 12(1). 10.1038/s41467-021-26751-5

Hochstein, S., & Ahissar, M. (2002). View from the top: Hierarchies and reverse hierarchies in the visual system. In Neuron (Vol. 36, Issue 5). 10.1016/S0896-6273(02)01091-7

Hung, C. P., Kreiman, G., Poggio, T., & DiCarlo, J. J. (2005). Fast readout of object identity from macaque inferior temporal cortex. Science, 310(5749). 10.1126/science.1117593

Hussar, C. R., & Pasternak, T. (2009). Flexibility of Sensory Representations in Prefrontal Cortex Depends on Cell Type. Neuron, 64(5). 10.1016/j.neuron.2009.11.018

Itti, L., & Koch, C. (2000). A saliency-based search mechanism for overt and covert shifts of visual attention. Vision Research, 40(10–12). 10.1016/S0042-6989(99)00163-7

Jiang, L. P., & Rao, R. P. N. (2024). Dynamic predictive coding: A model of hierarchical sequence learning and prediction in the neocortex. PLoS Computational Biology, 20(2). 10.1371/journal.pcbi.1011801

Kingma, D. P., & Welling, M. (2014). Auto-encoding variational bayes. 2nd International Conference on Learning Representations, ICLR 2014 - Conference Track Proceedings. 10.61603/ceas.v2i1.33

Koch, C., & Ullman, S. (1985). Shifts in selective visual attention: Towards the underlying neural circuitry. Human Neurobiology, 4(4). 10.1007/978-94-009-3833-5_5

Kriegeskorte, N., Mur, M., Ruff, D. A., Kiani, R., Bodurka, J., Esteky, H., Tanaka, K., & Bandettini, P. A. (2008). Matching Categorical Object Representations in Inferior Temporal Cortex of Man and Monkey. Neuron, 60(6). 10.1016/j.neuron.2008.10.043

Kummerer, M., Wallis, T. S. A., Gatys, L. A., & Bethge, M. (2017). Understanding Low- and High-Level Contributions to Fixation Prediction. Proceedings of the IEEE International Conference on Computer Vision, 2017-October. 10.1109/ICCV.2017.513

Lawrence, S. J. D., van Mourik, T., Kok, P., Koopmans, P. J., Norris, D. G., & de Lange, F. P. (2018). Laminar Organization of Working Memory Signals in Human Visual Cortex. Current Biology, 28(21). 10.1016/j.cub.2018.08.043

Li, F. F., VanRullen, R., Koch, C., & Perona, P. (2002). Rapid natural scene categorization in the near absence of attention. Proceedings of the National Academy of Sciences of the United States of America, 99(14). 10.1073/pnas.092277599

Locatello, F., Weissenborn, D., Unterthiner, T., Mahendran, A., Heigold, G., Uszkoreit, J., Dosovitskiy, A., & Kipf, T. (2020). Object-centric learning with slot attention. Advances in Neural Information Processing Systems, 2020-December.

Marchette, S. A., Vass, L. K., Ryan, J., & Epstein, R. A. (2015). Outside looking in: Landmark generalization in the human navigational system. Journal of Neuroscience, 35(44). 10.1523/JNEUROSCI.2270-15.2015

Marino, J. (2021). Predictive coding, variational autoencoders, and biological connections. In Neural Computation (Vol. 34, Issue 1). 10.1162/neco_a_01458

Martinez-Trujillo, J. (2022). Visual Attention in the Prefrontal Cortex. In Annual review of vision science (Vol. 8). 10.1146/annurev-vision-100720-031711

McInnes, L., Healy, J., Saul, N., & Großberger, L. (2018). UMAP: Uniform Manifold Approximation and Projection. Journal of Open Source Software, 3(29). 10.21105/joss.00861

Mégevand, P., Groppe, D. M., Goldfinger, M. S., Hwang, S. T., Kingsley, P. B., Davidesco, I., & Mehta, A. D. (2014). Seeing scenes: Topographic visual hallucinations evoked by direct electrical stimulation of the parahippocampal place area. Journal of Neuroscience, 34(16). 10.1523/JNEUROSCI.5202-13.2014

Mendoza-Halliday, D., & Martinez-Trujillo, J. C. (2017). Neuronal population coding of perceived and memorized visual features in the lateral prefrontal cortex. Nature Communications, 8. 10.1038/ncomms15471

Motoyoshi, I., Nishida, S., Sharan, L., & Adelson, E. H. (2007). Image statistics and the perception of surface qualities. Nature, 447(7141), 206– 209. 10.1038/NATURE05724;KWRD=SCIENCE

Nagy, D. G., Török, B., & Orbán, G. (2020). Optimal forgetting: Semantic compression of episodic memories. PLoS Computational Biology, 16(10). 10.1371/journal.pcbi.1008367

Navon, D. (1977). Forest before trees: The precedence of global features in visual perception. Cognitive Psychology, 9(3). 10.1016/0010-0285(77)90012-3

Oliva, A. (2005). Gist of the Scene. In Neurobiology of Attention. 10.1016/B978-012375731-9/50045-8

Oliva, A., & Torralba, A. (2006). Chapter 2 Building the gist of a scene: the role of global image features in recognition. In Progress in Brain Research: Vol. 155 B. 10.1016/S0079-6123(06)55002-2

Park, J., & Park, S. (2017). Conjoint representation of texture ensemble and location in the parahippocampal place area. Journal of Neurophysiology, 117(4). 10.1152/jn.00338.2016

Park, S., Brady, T. F., Greene, M. R., & Oliva, A. (2011). Disentangling scene content from spatial boundary: Complementary roles for the parahippocampal place area and lateral occipital complex in representing real-world scenes. Journal of Neuroscience, 31(4). 10.1523/JNEUROSCI.3885-10.2011

Peelen, M. V., Berlot, E., & de Lange, F. P. (2024). Predictive processing of scenes and objects. In Nature Reviews Psychology (Vol. 3, Issue 1). 10.1038/s44159-023-00254-0

Raju, R. V., Guntupalli, J. S., Zhou, G., Wendelken, C., Lázaro-Gredilla, M., & George, D. (2024). Space is a latent sequence: A theory of the hippocampus. Science Advances, 10(31), 31. 10.1126/SCIADV.ADM8470/SUPPL_FILE/SCIADV.ADM8470_SM.PDF

Rensink, R. A., O’Regan, J. K., & Clark, J. J. (1997). To see or not to see: The Need for Attention to Perceive Changes in Scenes. Psychological Science, 8(5). 10.1111/j.1467-9280.1997.tb00427.x

Reynolds, J. H., & Chelazzi, L. (2004). Attentional modulation of visual processing. In Annual Review of Neuroscience (Vol. 27). 10.1146/annurev.neuro.26.041002.131039

Rolls, E. T. (2024). Two what, two where, visual cortical streams in humans. Neuroscience & Biobehavioral Reviews, 160, 105650. 10.1016/J.NEUBIOREV.2024.105650

Rosenholtz, R., Huang, J., Raj, A., Balas, B. J., & Ilie, L. (2012). A summary statistic representation in peripheral vision explains visual search. Journal of Vision, 12(4). 10.1167/12.4.14

Rust, N. C., & DiCarlo, J. J. (2010). Selectivity and tolerance (“invariance”) both increase as visual information propagates from cortical area V4 to IT. Journal of Neuroscience, 30(39). 10.1523/JNEUROSCI.0179-10.2010

Scholte, H. S., Jolij, J., Fahrenfort, J. J., & Lamme, V. A. F. (2008). Feedforward and recurrent processing in scene segmentation: Electroencephalography and functional magnetic resonance imaging. Journal of Cognitive Neuroscience, 20(11). 10.1162/jocn.2008.20142

Seijdel, N., Loke, J., van de Klundert, R., van der Meer, M., Quispel, E., van Gaal, S., de Haan, E. H. F., & Scholte, H. S. (2021). On the necessity of recurrent processing during object recognition: It depends on the need for scene segmentation. Journal of Neuroscience, 41(29). 10.1523/JNEUROSCI.2851-20.2021

Silson, E. H., Chan, A. W. Y., Reynolds, R. C., Kravitz, D. J., & Baker, C. I. (2015). A retinotopic basis for the division of high-level scene processing between lateral and ventral human occipitotemporal cortex. Journal of Neuroscience, 35(34). 10.1523/JNEUROSCI.0137-15.2015

Silson, E. H., Groen, I. I. A., Kravitz, D. J., & Baker, C. I. (2016). Evaluating the correspondence between face-, scene-, and object-selectivity and retinotopic organization within lateral occipitotemporal cortex. Journal of Vision, 16(6). 10.1167/16.6.14

Silson, E. H., Steel, A. D., & Baker, C. I. (2016). Scene-selectivity and retinotopy in medial parietal cortex. Frontiers in Human Neuroscience, 10. 10.3389/fnhum.2016.00412

Simons, D. J., & Levin, D. T. (1997). Change blindness. Trends in Cognitive Sciences, 1(7), 261–267. 10.1016/S1364-6613(97)01080-2

Stachenfeld, K. L., Botvinick, M. M., & Gershman, S. J. (2017). The hippocampus as a predictive map. Nature Neuroscience, 20(11). 10.1038/nn.4650

Summerfield, C., & De Lange, F. P. (2014). Expectation in perceptual decision making: Neural and computational mechanisms. In Nature Reviews Neuroscience (Vol. 15, Issue 11). 10.1038/nrn3838

Tanaka, K. (1996). Inferotemporal cortex and object vision. In Annual Review of Neuroscience (Vol. 19). 10.1146/annurev.ne.19.030196.000545

Thorpe, S., Fize, D., & Marlot, C. (1996). Speed of processing in the human visual system. Nature, 381(6582). 10.1038/381520a0

Vafaii, H., Yates, J. L., & Butts, D. (2023). Hierarchical VAEs provide a normative account of motion processing in the primate brain. BioRxiv.

Wardak, C., Ibos, G., Duhamel, J. R., & Olivier, E. (2006). Contribution of the monkey frontal eye field to covert visual attention. Journal of Neuroscience, 26(16). 10.1523/JNEUROSCI.3336-05.2006

Watt, R. J. (1987). Scanning from coarse to fine spatial scales in the human visual system after the onset of a stimulus. Journal of the Optical Society of America A, 4(10). 10.1364/josaa.4.002006

Whitney, D., & Yamanashi Leib, A. (2018). Ensemble Perception. In Annual Review of Psychology (Vol. 69). 10.1146/annurev-psych-010416-044232

Wolfe, J. M. (1994). Guided Search 2.0 A revised model of visual search. Psychonomic Bulletin & Review, 1(2). 10.3758/BF03200774

Wolfe, J. M. (2021). Guided Search 6.0: An updated model of visual search. In Psychonomic Bulletin and Review (Vol. 28, Issue 4). 10.3758/s13423-020-01859-9

Wolfe, J. M., & Horowitz, T. S. (2017). Five factors that guide attention in visual search. Nature Human Behaviour, 1(3). 10.1038/s41562-017-0058

Wolfe, J. M., Võ, M. L. H., Evans, K. K., & Greene, M. R. (2011). Visual search in scenes involves selective and nonselective pathways. In Trends in Cognitive Sciences (Vol. 15, Issue 2). 10.1016/j.tics.2010.12.001

Yamins, D. L. K., Hong, H., Cadieu, C. F., Solomon, E. A., Seibert, D., & DiCarlo, J. J. (2014). Performance-optimized hierarchical models predict neural responses in higher visual cortex. Proceedings of the National Academy of Sciences of the United States of America, 111(23). 10.1073/pnas.1403112111

Yao, M., Wen, B., Yang, M., Guo, J., Jiang, H., Feng, C., Cao, Y., He, H., & Chang, L. (2023). High-dimensional topographic organization of visual features in the primate temporal lobe. Nature Communications, 14(1). 10.1038/s41467-023-41584-0

Zhuang, C., Yan, S., Nayebi, A., Schrimpf, M., Frank, M. C., DiCarlo, J. J., & Yamins, D. L. K. (2021). Unsupervised neural network models of the ventral visual stream. Proceedings of the National Academy of Sciences of the United States of America, 118(3). 10.1073/pnas.2014196118

